# Volumetric Differences of Thalamic Nuclei are Associated with Post-Trauma Psychopathology

**DOI:** 10.1101/2025.07.24.666644

**Authors:** Nick Steele, Ahmed Hussain, C. Lexi Baird, Courtney C. Haswell, Delin Sun, Leonel Rangel-Jimenez, Chadi G. Abdallah, Michael Angstadt, Geoffrey May, Hannah Berg, Jennifer U. Blackford, Josh M. Cisler, Judith K. Daniels, Nicholas D. Davenport, Richard J. Davidson, Maria Densmore, Seth G. Disner, Wissam El-Hage, Amit Etkin, Negar Fani, Jessie L. Frijling, Evan M. Gordon, Daniel W. Grupe, Ryan J. Herringa, Anna R. Hudson, Neda Jahanshad, Tanja Jovanovic, Anthony King, Saskia B.J. Koch, Ruth Lanius, Amit Lazarov, Gen Li, Israel Liberzon, Shmuel Lissek, Guangming Lu, Antje Manthey, Adi Maron-Katz, Laura Nawijn, Steven M. Nelson, Yuval Neria, Richard W.J. Neufeld, Jack B. Nitschke, Bunmi O. Olatunji, Miranda Olff, Matthew Peverill, Yann Quidé, Orren Ravid, Gopalkumar Rakesh, Kerry Ressler, Marisa Ross, Kelly Sambrook, Anika Sierk, Scott R. Sponheim, Jennifer Stevens, Benjamin Suarez-Jimenez, Jean Théberge, Sanne J.H. van Rooij, Mirjam van Zuiden, Dick J. Veltman, Robert R.J.M. Vermeiren, Henrik Walter, Li Wang, Xi Zhu, Ye Zhu, Sigal Zilcha-Mano, Christine Larson, Terri A. deRoon-Cassini, Carissa W. Tomas, Jacklynn M. Fitzgerald, Andrew S. Cotton, Erin N. O’Leary, Hong Xie, Xin Wang, Emily L. Dennis, David F. Tate, David X. Cifu, William C. Walker, Elisabeth A. Wilde, Paul M. Thompson, Rajendra A. Morey

## Abstract

Previous investigations of whole thalamus and thalamic nuclei volumes in post-trauma psychopathology have been sparse, limited in scope, and yielded inconsistent results. To address this, volumetric estimates of whole thalamus and thalamic nuclei were obtained from structural brain MRI scans from 2,058 participants across 20 worldwide sites in the ENIGMA PTSD working group. Thalamic volumes were compared between trauma-exposed participants with posttraumatic stress disorder (PTSD) (*n*=238), major depressive disorder (MDD) (*n*=184), comorbid PTSD+MDD (*n*=618), and trauma-exposed control participants (*n*=1,018). PTSD and MDD symptom severity, PTSD symptom clusters, and childhood trauma were similarly examined for associations with thalamic volume. Participants with PTSD only compared to controls had smaller thalamic nuclei volumes in sensorimotor nuclei, including the parafascicular (Pf), ventral anterior magnocellular (VAmc), medial pulvinar (PuM), and anterior pulvinar (PuA) nuclei of the thalamus. MDD only and comorbid PTSD+MDD participants exhibited smaller mediodorsal thalamus volumes compared to controls. Overall PTSD and MDD symptom severity negatively correlated with the volume of the mediodorsal thalamus. A significant interaction between PTSD and MDD severity was found, such that MDD severity was positively associated with thalamic volume only among individuals with high PTSD severity. Avoidance and hyperarousal symptoms of PTSD were positively associated with thalamic volume, while re-experiencing and negative mood/cognition symptoms were negatively associated with thalamic volume. Childhood physical and emotional abuse were positively and negatively associated with thalamic volume, respectively. Whole thalamus volume and volumes of the sensorimotor and limbic thalamus may play an important role in the development of PTSD and MDD in the aftermath of trauma exposure. The interaction between PTSD and MDD symptoms and contrasting effects across PTSD symptom clusters and types of childhood adversity suggests multiple neurobiological mechanisms are involved in shaping thalamic volume post-trauma.

## Introduction

Upwards of 85% of individuals experience a traumatic event at some point in their lives (1). Post-trauma psychopathology develops in a sizable minority of those exposed to trauma. For instance, posttraumatic stress disorder (PTSD) has a lifetime prevalence of 6.8% in the United States (2), and in many cases major depressive disorder (MDD) occurs independently of PTSD following a traumatic event (3,4). While the prevalence of PTSD following trauma exposure is greater than MDD in the near-term, they have similar long-term prevalence rates (5). Comorbid PTSD and MDD, occurring in around 50% of PTSD patients, typically results in more severe symptomatology and more treatment resistant illness (4,6,7). These challenges have led researchers to view these conditions not as completely independent disorders but as related aspects of a wider construct of post-trauma psychopathology (8).

Dysfunction of the hippocampus, amygdala, and ventromedial prefrontal cortex (vmPFC) resulting in altered fear learning and processing of affective information has been extensively demonstrated in post-trauma disorders (9,10). However, the thalamus has received less attention despite evidence of altered functional properties within post-trauma psychiatric groups (11–13). The thalamus is a key node in perceptual, cognitive, and affective processing circuits. It receives input from nearly every sensory organ system and reciprocally connects with virtually the entire brain, making it crucial for cortico-cortical communication, cortical arousal, modulating perceptual information, and integrating affective and memory-related information relevant to behavioral goals (14,15). Volumetric reductions or enlargements of the thalamus and thalamic nuclei have been linked to numerous psychopathologies including schizophrenia (16), obsessive- compulsive disorder (17), and bipolar disorder (18).

Thalamic volume has been consistently associated with trauma and post-trauma symptoms, yet the directionality of findings has been mixed. Individuals who develop PTSD or experience more severe PTSD symptoms following a traumatic event have lower volume of the thalamus compared to trauma-exposed individuals who do not develop PTSD (19–22).

Conversely, thalamic nuclei volumes have been positively correlated with the overall severity of PTSD symptoms (21) and both positively and negatively correlated with PTSD hyperarousal and re-experiencing symptoms (21,23,24). The relationship between depressive symptoms or MDD diagnosis and thalamic volume following trauma remains relatively unexplored. MDD diagnosis in non-trauma-exposed samples has been associated with lower whole thalamus and thalamic nuclei volumes (22,25,26). Among veterans with either MDD or PTSD, a greater number of suicidal behaviors – a sign of severe symptomatology – related to higher whole thalamus volume (27). More severe childhood trauma is associated with lower volume of the thalamus among individuals with PTSD (19,20). Individuals with a history of childhood trauma are also at a greater risk of developing psychopathology following trauma in adulthood (28). Thus, understanding the relationships between childhood trauma, post-trauma psychiatric symptoms, and thalamic volume is crucial to our overall understanding of how trauma impacts the brain.

The aforementioned studies had relatively limited sample sizes, ranging from *n*=32 to *n*=152. Research suggests associations between morphometric brain measures and psychiatric symptoms become robust and reproducible as sample size reaches thousands of participants (29). Smaller sample sizes result in more homogeneous samples, limiting the ability to generalize to a wider population. Small samples also lead to greater sampling variability across studies which increases sign-error and reduces reproducibility of results. A prior ENIGMA PTSD study investigated subcortical brain volumes in PTSD with a large sample (*n*=1,868) and found smaller thalamic volume in PTSD, but the result did not survive adjustment for intracranial volume or multiple comparison correction (30). However, several limitations of the study, including using meta-analysis rather than mega-analysis and only examining whole thalamus volume and not thalamic nuclei volumes, restrict the conclusions that can be drawn. To address previous limitations, we analyzed structural brain MRI scans from 2,058 participants by aggregating data from 20 worldwide sites in the ENIGMA PTSD working group. Volumes of the left and right thalamus and 25 individual thalamic nuclei per hemisphere were compared between trauma-exposed control participants and participants with post-trauma psychopathology, which include PTSD only, MDD only, and comorbid PTSD+MDD. Based on previous findings, we hypothesized volumetric differences of the ventral motor nuclei and mediodorsal (MD) thalamus in relation to psychiatric diagnosis, symptom severity, and childhood trauma.

## Methods

### Sample

Clinical and demographic data on the sample from the ENIGMA PTSD working group are shown in *Table 1*. T1-weighted brain MRI scans were collected from 20 sites (see *Supplementary Table 1* for demographics by site and *Supplementary Table 2* for inclusion and exclusion criteria by site). After removal of statistical outliers and participants with incomplete data, 2,058 participants remained in the final analysis, including 1,040 participants with post-trauma psychopathology and 1,018 control participants (95.8% trauma-exposed). All study procedures were approved by local institutional review boards (IRB), and all participants provided written informed consent. Secondary data analysis was deemed exempt by the Duke University Medical Center IRB.

**Table 1:** Demographic and clinical data. Means reported with standard deviations in parentheses. Symptom severity scores have been normalized between zero and one (see below). MDD data not available for ∼20% of CTQ subset. CTQ = Childhood trauma questionnaire.

|  | Controls | PTSD Only | MDD Only | Comorbid PTSD+MDD | Statistic | Significant Pairwise Comparisons |
| --- | --- | --- | --- | --- | --- | --- |
| *N* | 1,018 | 238 | 184 | 618 | - | - |
| Age | 38.83 (11.01) | 38.99 (10.92) | 38.38 (10.51) | 38.72 (10.54) | $F = 0.08, p = 0.969$ | - |
| Age Range | 18-72 | 18-67 | 19-62 | 18-68 | - | - |
| Female (%) | 25.34 | 42.02 | 34.78 | 40.45 | $\chi^2 = 50.17, p < 0.001$ | Control-Comorbid; Control-MDD Only; Control-PTSD Only |
| PTSD Severity | 0.11 (0.11) | 0.46 (0.13) | 0.26 (0.12) | 0.58 (0.14) | $F = 1931.08, p < 0.001$ | All pairwise comparisons significant |
| MDD Severity | 0.11 (0.10) | 0.21 (0.09) | 0.45 (0.12) | 0.52 (0.16) | $F = 1715.71, p < 0.001$ | All pairwise comparisons significant |
| **PTSD Symptoms** |  |  |  |  |  |  |
| *N* | 285 | 115 | 55 | 264 | - | - |
| Re-experiencing | 0.09 (0.13) | 0.42 (0.18) | 0.25 (0.20) | 0.55 (0.19) | $F = 465.62, p < 0.001$ | All pairwise comparisons significant |
| Avoidance | 0.10 (0.18) | 0.53 (0.22) | 0.24 (0.23) | 0.61 (0.22) | $F = 316.65, p < 0.001$ | All pairwise comparisons significant |
| Negative Mood/Cognition | 0.04 (0.08) | 0.37 (0.15) | 0.14 (0.15) | 0.50 (0.19) | $F = 491.52, p < 0.001$ | All pairwise comparisons significant |
| Hyperarousal | 0.11 (0.14) | 0.43 (0.18) | 0.29 (0.19) | 0.56 (0.16) | $F = 505.93, p < 0.001$ | All pairwise comparisons significant |
| **CTQ Severity** |  |  |  |  |  |  |
| *N* | 270 | 75 | 54 | 156 | - | - |
| CTQ Total | 37.37 (13.78) | 56.49 (24.14) | 40.96 (16.09) | 62.77 (23.92) | $F = 53.15, p < 0.001$ | Control-Comorbid; MDD Only-Comorbid; PTSD Only-Control |
| Emotional Abuse | 7.91 (3.94) | 13.21 (6.36) | 8.65 (4.53) | 15.35 (6.59) | $F = 58.77, p < 0.001$ | Control-Comorbid; MDD Only-Comorbid; PTSD Only-Comorbid; PTSD Only-Control |
| Physical Abuse | 7.42 (3.59) | 9.92 (5.14) | 8.31 (4.32) | 10.84 (5.61) | $F = 17.55, p < 0.001$ | Control-Comorbid; PTSD Only-Control |
| Sexual Abuse | 6.58 (3.98) | 10.72 (6.96) | 6.96 (4.67) | 11.74 (6.95) | $F = 21.87, p < 0.001$ | Control-Comorbid; PTSD Only-Control |
| Emotional Neglect | 8.84 (4.24) | 13.01 (6.07) | 9.48 (4.38) | 14.72 (5.97) | $F = 36.97, p < 0.001$ | Control-Comorbid; PTSD Only-Control |
| Physical Neglect | 6.62 (2.52) | 9.63 (4.50) | 7.56 (3.14) | 10.12 (4.56) | $F = 33.82, p < 0.001$ | Control-Comorbid; MDD Only-Comorbid; PTSD Only-Control |

### Assessment and Harmonization of Clinical Data Across Sites

Clinical diagnosis of PTSD was determined using a specific assessment tool at each site based on DSM-IV or DSM-5 criteria, which differed by site. Test-specific cut-off scores were used to determine MDD status. Severity of psychiatric symptoms were measured using a range of questionnaires and clinical interviews across the 20 sites (see *Supplementary Table 1*). To allow for comparison of these different measures with each other, PTSD, MDD, and symptom cluster scores were normalized between zero and one. This was done by subtracting the minimum possible score from each participant’s score then dividing by the maximum possible score minus the minimum possible score, resulting in a severity ranging from zero to one {severity∈R∣0<x<1}. No harmonization was performed for childhood trauma scores because all sites used the same test (Childhood Trauma Questionnaire [CTQ]).

### Image Acquisition and Processing

T1-weighted brain MRI scans were acquired from each site (see *Supplementary Table 3* for scanning parameters of each site) and were processed via FreeSurfer (version 7.1.1). The *SegmentThalamicNuclei* pipeline within FreeSurfer (31) was then applied to segment the thalamus into 25 distinct nuclei in each hemisphere. Volumetric estimates were obtained for the left and right whole thalamus and for each of the 50 thalamic nuclei (*Figure 1*). The thalamus can be broadly divided into six subregions: the posterior, ventral, anterior, lateral, medial, and intralaminar regions. An alternative classification scheme divides the thalamus by nuclei function and recognizes sensorimotor nuclei, limbic nuclei, and association nuclei involved in both sensorimotor and limbic processes (32).

**Figure 1:**
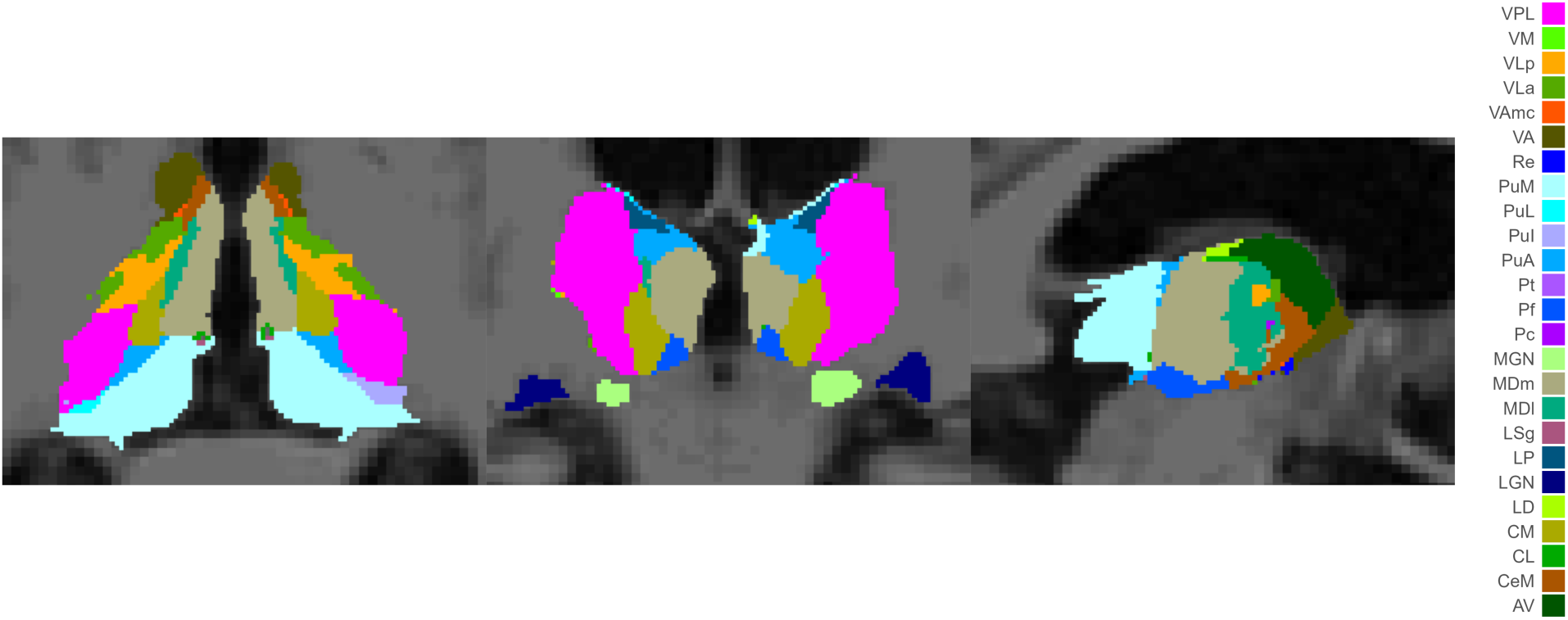
Axial (left), coronal (middle), and sagittal (right) views of an example thalamic segmentation in subject-space with thalamic nuclei color coded.

Nuclei in the segmentation include sensorimotor nuclei in the posterior subregion [lateral geniculate nucleus (LGN); medial geniculate nucleus (MGN); limitans-suprageniculate (LSg); and anterior, inferior, lateral and medial pulvinar (PuA, PuI, PuL, PuM)] and ventral subregion [ventral posterolateral (VPL), ventromedial (VM), ventral lateral anterior (VLa), ventral lateral posterior (VLp), ventral anterior (VA), and ventral anterior magnocellular (VAmc)]; limbic nuclei in the medial subregion [mediodorsal medial (MDm), mediodorsal lateral (MDl), paratenial (Pt), and reuniens (Re)], intralaminar subregion [central medial (CeM)], and anterior subregion [anteroventral (AV)]; and association nuclei in the intralaminar subregion [central lateral (CL), paracentral (Pc), centromedian (CM), parafascicular (Pf)] and lateral subregion [lateral posterior (LP) and lateral dorsal (LD)].

To address the problem of potential missegmentations, statistical outliers were removed following standard ENIGMA quality control procedures for structural segmentations.

Segmentations with any thalamic hemisphere or nuclei that exceeded ±2.698 standard deviations (33) from the sample mean for that region were removed completely from statistical analysis (Controls *n*=234, PTSD only *n*=60, MDD only *n*=31, comorbid PTSD+MDD *n*=111).

### Volume Harmonization Across Sites

To adjust for the effects of site on thalamic volume, the *neuroHarmonize* package in python (version 3.9.19) was used to adjust for variability related to site while preserving variability related to the variables of interest (Pomponio et al., 2020). The *neuroHarmonize* package implements the *ComBat-GAM* algorithm based on an empirical Bayes framework. For sites that collected data from multiple MRI scanners, each scanner was treated as a separate site to adjust for scanner-related variability in thalamic volume. All harmonization models included sex, age, and intracranial volume (ICV) as covariates, with age being set as a smooth term to account for non-linear effects, in addition to the clinical variables of interest. The site harmonized thalamic volumes were then entered into linear models to test for significant associations between thalamic volume and the clinical variables of interest.

### Statistical Analysis

The effects of psychopathology on whole thalamus and thalamic nuclei volume were examined. All models included sex, age, age^2^, and ICV as covariates of no interest. Age^2^ was included due to the large age range of our sample and the known non-linear effects of age on brain volume (35,36).

An analysis of covariance (ANCOVA) was used to test for group differences between four diagnostic groups including participants with 1) PTSD only, 2) MDD only, 3) comorbid PTSD+MDD, and 4) controls without PTSD or MDD (95.8% trauma-exposed) (Equation 1). Post-hoc pairwise comparisons were performed on significant ANCOVA results. Cohen’s *d* was used as a measure of effect size. Tests for the 50 thalamic nuclei were corrected for multiple comparisons using the Benjamini-Hochberg procedure with a false discovery rate (FDR) of *q=*0.05 (37), and *p*-values of post-hoc pairwise comparisons were corrected for the six group comparisons using the Tukey Honestly Significant Difference (HSD) method.

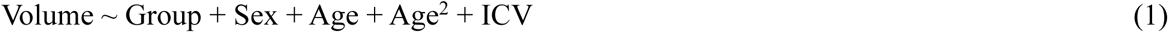

PTSD and MDD symptom severity was tested for associations with thalamic volume (Equations 2-3). The interaction between PTSD and MDD severity was examined as well (Equation 4).

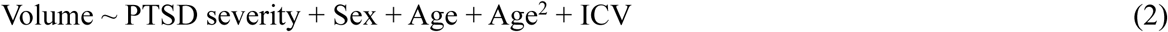

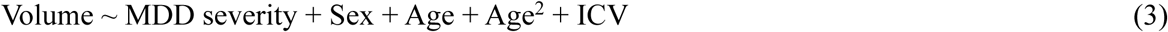

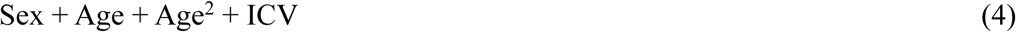

Specific psychiatric symptoms were examined by testing the association between DSM-5 symptom clusters for PTSD (criteria B, C, D, and E) and thalamic volume (Equation 5). These symptom clusters refer to re-experiencing, avoidance, negative mood/cognition, and hyperarousal symptoms, respectively. Each symptom cluster was analyzed while controlling for the other symptoms to better understand the unique contributions of each symptom cluster.

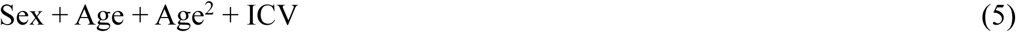

Finally, childhood trauma was measured via the CTQ (Bernstein et al., 1994). Total CTQ scores and CTQ subscores – emotional abuse (EA), physical abuse (PA), sexual abuse (SA), emotional neglect (EN), and physical neglect (PN) – were tested for associations with thalamic volume (Equation 6-7).

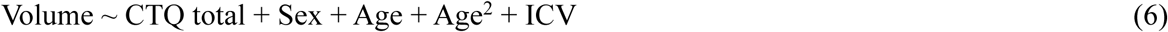

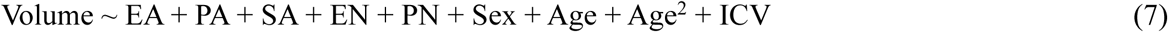

Dominance analysis was used to determine which model terms contributed the most to differences in thalamic volume using the *domir* package in R (39). Model terms of interest were ranked according to which term was assigned the highest proportion of model fit, called general dominance. Model terms can generally dominate, conditionally dominate, or completely dominate other model terms. Conditional dominance is more stringent than general dominance and occurs when a model term’s average additional contribution to model fit is greater than that of another model term across all model sizes. Complete dominance is the most stringent form of dominance, which occurs when a larger percentage of model fit is attributed to a model term over another term in every possible sub-model.

## Results

### Psychiatric Diagnosis and Thalamic Volume

Statistically significant ANCOVA (Equation 1) results are available in *Table 2* and full statistical results are available in *Supplementary Table 4*. Several thalamic nuclei showed significant volumetric differences between diagnostic groups (*Figure S1*).

**Table 2:**
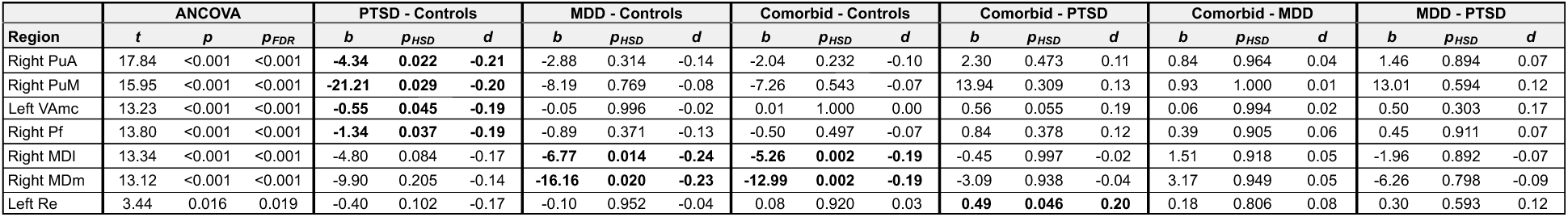
Significant results from pairwise comparisons between diagnostic groups. The p-values for ANCOVA results have been FDR-corrected for the 50 thalamic nuclei, while p-values for pairwise comparisons have been corrected using the Tukey HSD method for the six group comparisons. Bolded values represent significant pairwise results.

| Region | ANCOVA |  |  | PTSD - Controls |  |  | MDD - Controls |  |  | Comorbid - Controls |  |  | Comorbid - PTSD |  |  | Comorbid - MDD |  |  | MDD - PTSD |  |  |
| --- | --- | --- | --- | --- | --- | --- | --- | --- | --- | --- | --- | --- | --- | --- | --- | --- | --- | --- | --- | --- | --- |
|  | *t* | *p* | *pFDR* | *b* | *pHSD* | *d* | *b* | *pHSD* | *d* | *b* | *pHSD* | *d* | *b* | *pHSD* | *d* | *b* | *pHSD* | *d* | *b* | *pHSD* | *d* |
| Right PuA | 17.84 | <0.001 | <0.001 | **-4.34** | **0.022** | **-0.21** | -2.88 | 0.314 | -0.14 | -2.04 | 0.232 | -0.10 | 2.30 | 0.473 | 0.11 | 0.84 | 0.964 | 0.04 | 1.46 | 0.894 | 0.07 |
| Right PuM | 15.95 | <0.001 | <0.001 | **-21.21** | **0.029** | **-0.20** | -8.19 | 0.769 | -0.08 | -7.26 | 0.543 | -0.07 | 13.94 | 0.309 | 0.13 | 0.93 | 1.000 | 0.01 | 13.01 | 0.594 | 0.12 |
| Left VAmc | 13.23 | <0.001 | <0.001 | **-0.55** | **0.045** | **-0.19** | -0.05 | 0.996 | -0.02 | 0.01 | 1.000 | 0.00 | 0.56 | 0.055 | 0.19 | 0.06 | 0.994 | 0.02 | 0.50 | 0.303 | 0.17 |
| Right Pf | 13.80 | <0.001 | <0.001 | **-1.34** | **0.037** | **-0.19** | -0.89 | 0.371 | -0.13 | -0.50 | 0.497 | -0.07 | 0.84 | 0.378 | 0.12 | 0.39 | 0.905 | 0.06 | 0.45 | 0.911 | 0.07 |
| Right MDI | 13.34 | <0.001 | <0.001 | -4.80 | 0.084 | -0.17 | **-6.77** | **0.014** | **-0.24** | **-5.26** | **0.002** | **-0.19** | -0.45 | 0.997 | -0.02 | 1.51 | 0.918 | 0.05 | -1.96 | 0.892 | -0.07 |
| Right MDm | 13.12 | <0.001 | <0.001 | -9.90 | 0.205 | -0.14 | **-16.16** | **0.020** | **-0.23** | **-12.99** | **0.002** | **-0.19** | -3.09 | 0.938 | -0.04 | 3.17 | 0.949 | 0.05 | -6.26 | 0.798 | -0.09 |
| Left Re | 3.44 | 0.016 | 0.019 | -0.40 | 0.102 | -0.17 | -0.10 | 0.952 | -0.04 | 0.08 | 0.920 | 0.03 | **0.49** | **0.046** | **0.20** | 0.18 | 0.806 | 0.08 | 0.30 | 0.593 | 0.12 |

Among participants with PTSD only, smaller volume was found in the right PuA, right PuM, left VAmc, and right Pf nuclei compared to control participants. Among participants with MDD only, smaller volumes were found in the right MDl and right MDm nuclei compared to control participants.

Participants with comorbid PTSD+MDD displayed smaller volumes of the right MDl and the right MDm nuclei compared to control participants. Additionally, comorbidity was associated with larger volume of the left Re nucleus compared to participants with PTSD only.

### Symptom Severity and Thalamic Volume

Symptom severity was tested for associations with thalamic volume (Equation 2-4) across the entire sample (*n*=2,058). PTSD symptom severity was significantly negatively correlated with left thalamus (*b*=-98.0, *p*=0.022) and right thalamus (*b=*-114.91, *p*=0.014) volume. Volumes of the right PuA (*b*=-5.85, *pFDR*=0.038), right MDl (*b*=-0.967, *pFDR*=0.004), and right MDm (*b*=-25.45, *pFDR*=0.003) nuclei were also significantly negatively correlated with PTSD severity.

MDD symptom severity did not significant correlate with left or right thalamus volume (*p*>0.05). However, volume of the right MDl (*b*=-11.24, *pFDR*=0.002) and right MDm (*b*=-24.25, *pFDR*=0.012) nuclei were significantly negatively correlated with MDD severity.

A significant interaction between PTSD severity and MDD severity was found (Equation 4), displaying a positive beta value for the right thalamus (*b*=502.47, *p*=0.008). Simple Slopes analysis revealed that PTSD severity significantly negatively correlated with right thalamus volume only when MDD severity is below 0.30 (*Figure 2A*). A significant positive relationship between MDD severity and right thalamus volume is observed only when PTSD severity is above 0.66 (*Figure 2B*).

**Figure 2:**
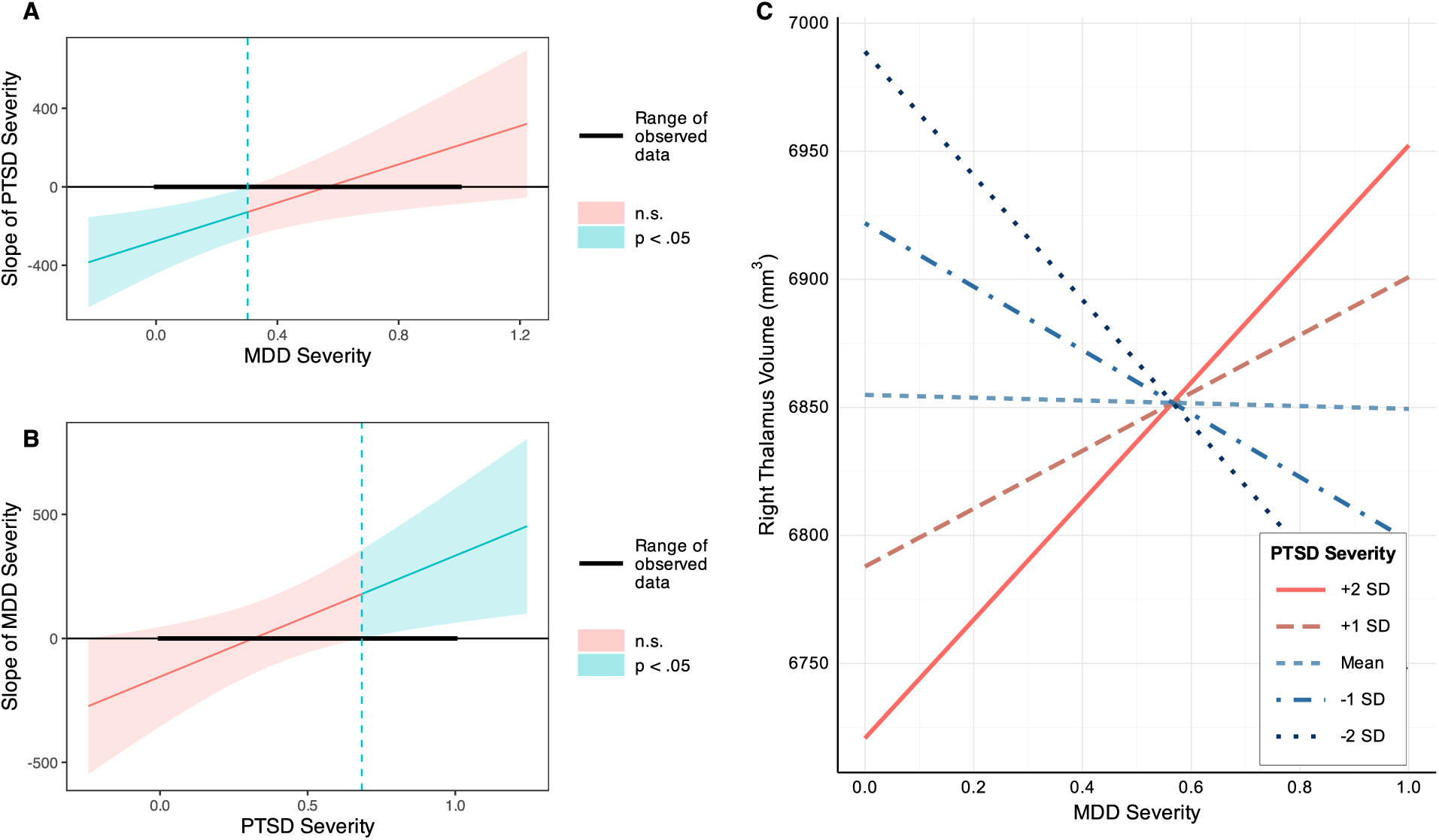
Interaction Between PTSD and MDD Severity. **(A)** The slope of PTSD severity is significantly negatively associated with right thalamus volume only when MDD severity is below 0.30. **(B)** The slope of MDD severity is significantly positively associated with right thalamus volume only when PTSD severity is above 0.66. **(C)** Right thalamus volume vs MDD symptom severity with PTSD severity standard deviations plotted. Among individuals with high PTSD symptoms, MDD severity is positively associated with right thalamus volume.

### PTSD Symptom Clusters and Thalamic Volume

Re-experiencing, avoidance, negative mood/cognition, and hyperarousal PTSD symptoms (DSM-5 criteria B, C, D, and E symptom clusters) were tested for associations with thalamic volume (Equation 5) across the entire sample with symptom-level data (*n*=719). The variance inflation factor (VIF) was calculated for each term to assess the impact of multicollinearity on model fit and stability (re-experiencing VIF=3.74, avoidance VIF=3.02, negative mood/cognition VIF=3.18, hyperarousal VIF=3.49, and MDD severity VIF=2.10). Further evaluation of multicollinearity was unwarranted given all VIF values were below five (40).

Re-experiencing symptoms were significantly negatively correlated with left thalamus (*b*=-258.6, *p*=0.043) and right thalamus (*b*=-315.4, *p*=0.015) volume. Negative mood/cognition symptoms were significantly negatively correlated with left thalamus (*b*=-287.4, *p*=0.019) volume. Conversely, avoidance symptoms were positively correlated with left thalamus (*b*=207.4, *p*=0.033) volume.

Regarding individual thalamic nuclei, re-experiencing symptoms were negatively correlated with volume in the right LP (*b*=-16.36, *pFDR*=0.044) nucleus. Hyperarousal symptoms were positively correlated with the right CL (*b*=6.52, *pFDR*=0.033), right LP (*b*=16.03, *pFDR*=0.033), and bilateral LD (left: *b*=6.86, *pFDR*=0.033; right: *b*=7.05, *pFDR*=0.033) nuclei.

For the left thalamus, dominance analysis revealed the largest proportion of model fit to be attributed to negative mood/cognition symptoms (3.52%), followed by re-experiencing (2.38%), avoidance (1.35%), hyperarousal (0.84%), and MDD severity (0.63%). For the right thalamus, negative mood/cognition symptoms accounted for 2.72% of model fit, followed by re-experiencing (2.70%), avoidance (1.29%), hyperarousal (0.90%), and MDD severity (0.57%).

### Childhood Trauma and Thalamic Volume

Childhood trauma was assessed using the CTQ. Total CTQ scores (Equation 6) and CTQ subscores (Equation 7) – EA, PA, SA, EN, and PN – were tested for associations with thalamic volume across the entire sample with CTQ data (*n*=694). To ensure multicollinearity was not leading to an unstable model fit, the VIF was calculated for each model term (EA VIF=4.30, PA VIF=2.08, SA VIF=1.76, EN VIF=3.95, and PN VIF=2.50). Further evaluation of multicollinearity was unwarranted given all VIF values were below five (40).

Total CTQ scores were not significantly associated with thalamic volume or thalamic nuclei volume (all *p*>0.05). Regarding the five CTQ subscores, PA was positively correlated with volume of the left thalamus (*b*=13.49, *p*=0.001), while EA was negatively correlated with left thalamus (*b*=-12.27, *p*=0.044) volume.

The left CeM (*b*=-0.40, *pFDR* = 0.024), left Re (*b*=-0.10, *pFDR* = 0.026), and left AV (*b*=- 0.72, *pFDR*=0.004) nuclei were significantly negatively correlated with EA scores. Volumetric analyses were rerun with PTSD and MDD diagnoses included in the model with consistent results. A summary of significant findings from all analyses can be seen in *Figure 3*.

**Figure 3:**
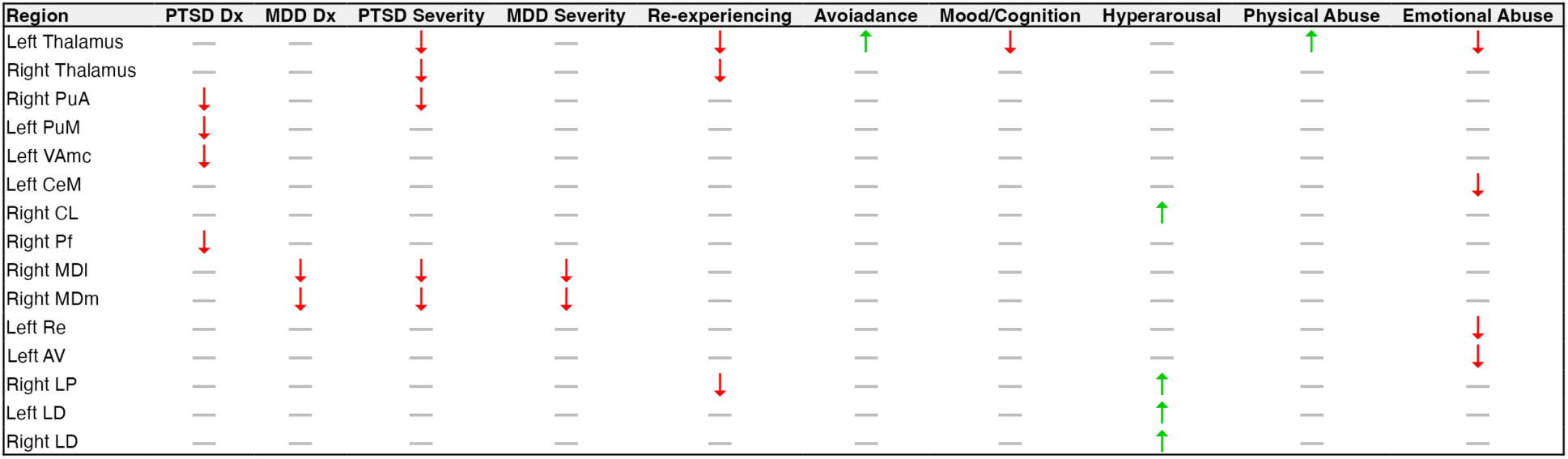
Summary of Results. Green up arrows represent findings of larger volume and red down arrows represent findings of smaller volume. The significant interaction term between PTSD and MDD severity is not shown.

Dominance analysis revealed the greatest proportion of model fit for predicting left thalamus volume to be attributed to EA (1.40%), followed by PA (0.85%), SA (0.65%), EN (0.37%), and PN (0.26%). For the right thalamus, SA ranked highest (1.04%), followed by PN (0.69%), EA (0.54%), PA (0.47%), and EN (0.30%).

## Discussion

The current investigation found lower volume among several nuclei spanning the ventral (VAmc), posterior (PuA, PuM), intralaminar (Pf), and medial (MDm, MDl) subregions of the thalamus in relation to diagnostic status of PTSD or MDD. Severity of PTSD symptoms negatively correlated with left and right whole thalamus volume, while both PTSD and MDD severity negatively correlated with mediodorsal thalamus volume. We found a significant negative correlation between PTSD re-experiencing and negative mood/cognition symptoms with thalamic volume, but there was a positive association between avoidance and hyperarousal symptoms with thalamic volume. Finally, childhood PA positively correlated with left thalamus volume, while childhood EA negatively correlated with left thalamus and thalamic nuclei volume.

### Differences in Thalamic Volume in Post-Trauma Psychopathology

Smaller thalamic nuclei volumes were observed in the PTSD only, MDD only, and comorbid PTSD+MDD groups compared to controls, and greater severity of PTSD and MDD symptoms was associated with smaller thalamus volume. However, we identified a significant interaction between PTSD and MDD severity, whereby high comorbidity was linked to larger thalamic volume, and the right Re nucleus displayed higher volume in the comorbid group compared to participants with PTSD only. These findings suggest that while mild to moderate post-trauma psychopathology is associated with reduced thalamic volume, severe symptomatology and comorbidity is associated with higher thalamic volume. Our results thus offer a nuanced explanation that partially reconciles previous findings of both smaller (19,21) and larger (24,27) thalamic volumes in trauma-related psychopathology. Prior investigations have reported reduced thalamic and thalamic subregion volumes in PTSD cases relative to trauma-exposed controls, yet patients with more severe symptoms exhibited greater thalamic volume compared to patients with mild symptoms (21). Aspects of severe symptomatology, such as suicidal behaviors, have also been associated with larger thalamic volumes (27). Together, these published results support our finding that severe symptomatology and comorbidity may be related to greater thalamic volume.

Severity of specific PTSD symptom clusters improved model fit to a greater extent than overall PTSD severity and MDD severity. Negative mood/cognition and re-experiencing symptoms contributed the most to model fit for predicting whole thalamus volume and were negatively associated with thalamic volume, while avoidance and hyperarousal symptoms were positively associated with thalamic volume. This evidence points to distinct neurobiological mechanisms acting upon the thalamus leading to differential relationships between distinct components of current symptomatology. Clinician assessment of patients with post-trauma psychiatric conditions and the development of individualized treatments may be improved by considering the degree of comorbidity and the severity of specific symptom profiles.

### Childhood Trauma and Thalamic Volume

Previous research reported an inverse association between childhood trauma and thalamic volume in PTSD (19). We found no significant association with total CTQ scores and thalamic volume but significant effects regarding CTQ subscores. Childhood abuse was significantly related to thalamic volume while childhood neglect was not. Moreover, childhood PA had a positive association with thalamic volume while childhood EA had a negative association. These findings align with evidence in the literature that different types of childhood adversity can lead to differential alterations in neurodevelopmental processes (41). Our results expand upon these ideas by demonstrating that the neurodevelopmental alterations implicated in distinct types of threat may be explained by specific brain changes to the thalamus.

### Volumetric Changes to the Sensorimotor Thalamus in PTSD

The PTSD only group exhibited lower volumes of the PuA, PuM, VAmc, and Pf nuclei compared to controls. While the severity of both PTSD and MDD symptoms was associated with mediodorsal thalamus volume, only PTSD severity related to PuA volume. Moreover, the LD, LP, and CL nuclei were positively associated with hyperarousal symptoms, and the LP nucleus was additionally negatively associated with re-experiencing symptoms. These nuclei are predominantly implicated in sensorimotor functions and have strong connections to sensory and/or motor cortices. VAmc and Pf nuclei contain strong projections with motor cortices, while the LD, LP, PuA, and PuM are considered visual association regions involved in visual or multisensory processing (42–44).

Volumetric changes in these regions may suggest a reduced ability to respond appropriately to incoming sensory information and to properly integrate information from various motor regions for adaptive motor responses. PTSD is characterized in part by somatomotor complaints and hyper- or hypo-responding to stimuli across sensory modalities leading to combinations of sensory sensitivity, low sensory registration, sensory avoidance, and sensation seeking (45). Altered sensory processing and multisensory integration has been theorized as a prominent component of the emotion dysregulation experienced in PTSD (46).

Clusters of symptoms related to sensorimotor responding, such as hyperarousal and avoidance symptoms (45), may explain volumetric changes to nuclei involved in sensory processing and the generation of motor responses.

### Volumetric Changes to the Limbic Thalamus in MDD and Childhood Emotional Abuse

We observed volumetric differences in the MDm, MDl, Re, CeM, and AV nuclei, which corroborates previous research (19,21). Both MDm and MDl nuclei volumes were found to be lower in MDD participants, comorbid PTSD+MDD participants, and negatively correlated with both PTSD and MDD symptom severity. The Re, CeM, and AV nuclei were negatively associated with childhood EA, and the Re nucleus additionally displayed larger volume in the comorbid PTSD+MDD group compared to the PTSD only group. These regions are considered limbic nuclei due to their strong connectivity to subcortical and cortical limbic structures. The MDm, Re, and AV nuclei have the strongest connectivity to the hippocampus of any thalamic nuclei (47,48). Dense cortical connections are found with prefrontal and retrosplenial regions, making these nuclei crucial for learning and memory, spatial processing, and goal-directed behavior (47–49). Given the dense connectivity of these nuclei to the hippocampus and vmPFC, in addition to the strong evidence of hippocampal and vmPFC dysfunction in post-trauma psychopathology (10), we hypothesize volumetric alterations to the limbic thalamus may play a role in impaired hippocampal-vmPFC communication post-trauma.

## Limitations and Strengths

This study has several important limitations, including a lack of information on comorbid conditions beyond PTSD and MDD, history of alcohol and substance use, and psychotropic medication usage. Nevertheless, the exclusion criteria employed by participating study sites likely reduced the potential impact of these confounding factors. Of the 20 contributing sites, 16 sites excluded individuals with a history of substance abuse, 17 sites excluded individuals with either any Axis I psychiatric disorder or a history of psychotic or manic symptoms, and 13 sites excluded individuals currently using any, or certain classes of, psychotropic medications. The exclusion of these individuals likely mitigated the influence of these potential confounds on our results. The sizable difference in the number of participants in each of the diagnostic groups is a limitation as well. Finally, a limitation of this study is the lack of standardized trauma load measures across all participating sites. While our control group is well-matched for binary trauma exposure (95.8% trauma-exposed), we cannot rule out that differences in trauma severity, chronicity, or type between diagnostic groups may have contributed to the observed thalamic volume differences.

Strengths of the present study are an order-of-magnitude larger sample size than any published study of thalamic nuclei volumes in post-trauma psychopathology. We examined volumetric differences in greater detail with 25 thalamic nuclei as compared to five or six regions used in previous studies. Employing a mega-analysis in a large multi-cohort consortium dataset enabled us to observe small effect sizes on thalamic nuclei. The clinical demography of our sample was diverse with respect to age, sex, clinical status, trauma type, race, and ancestry.

Therefore, our findings are likely to better generalize than previous studies with much smaller and homogenous samples and trauma types. Another strength is using *ComBat-GAM* harmonization to adjust for site- and scanner-effects. *ComBat-GAM* is currently recognized as one of the best methods for removing site- or scanner-related variability from neuroimaging datasets (50).

## Conclusions

Altered volumes of the limbic and sensorimotor thalamus in relation to post-trauma psychiatric diagnosis and symptom severity suggests that the thalamus may be associated with symptoms in these disorders. Contrasting effects of PTSD symptom clusters and types of childhood adversity suggests the possibility that multiple neurobiological mechanisms are involved in shaping thalamic volume post-trauma. Future research examining diagnostic status and changes in symptom severity over time is needed to understand whether altered thalamic volume constitutes a risk-factor or a consequence of psychiatric symptoms.

## Acknowledgements

The study was supported by ZonMw, the Netherlands organization for Health Research and Development (40-00812-98-10041), and by a grant from the Academic Medical Center Research Council (110614) both awarded to MO; the National Natural Science Foundation of China (No. U21A20364 and No. 31971020), the Key Project of the National Social Science Foundation of China (No. 20ZDA079), the Key Project of Research Base of Humanities and Social Sciences of Ministry of Education (No.16JJD190006), and the Scientific Foundation of Institute of Psychology, Chinese Academy of Sciences (No. E2CX4115CX) [to LW, YW]; NIMH K01MH118428 and NIMH R01MH131532 [to BSJ]; NARSAD 27040 and NIH K01MH122774 [to XZ]; RO1 MH111671, VISN6 MIRECC [to RM]; MH098212, MH071537, M01RR00039, UL1TR000454, HD071982; HD085850, Narsad Young Investigator, and MH101380 [to JS, KR, TJ]; Narsad Young Investigator [to SR]; MH101380 [to NF]; German Research Foundation grant to J. K. Daniels (numbers DA 1222/4-1 and WA 1539/8-2) [to JD, AS, AM, HW]; K99NS096116 [to ED]; HFP90-020 [to EW]; R01MH113574 [to IL]; R01 MH106574 [to CL, TD]; VA RR&D 1IK2RX000709 [to ND]; VA RR&D I01RX000622, CDMRP W81XWH-08–2–0038 [to SS]; VA RR&D 1K1RX002325, 1K2RX002922 [to SD]; 1R01MH110483, 1R21 MH098198 [to XW]; PHRC, Fondation Pierre Deniker and SFR FED4226 [to WEH]; VA CSR&D 1IK2CX001680, VISN17 Center of Excellence Pilot funding [to EG, GM, SN]; VA National Center for PTSD [to CA]; Dana Foundation (to Dr. Nitschke); the University of Wisconsin Institute for Clinical and Translational Research (to Dr. Emma Seppala); a National Science Foundation Graduate Research Fellowship (to Dr. Grupe); the National Institute of Mental Health (NIMH) R01- MH043454 and T32-MH018931 (to Dr. Davidson); the Canadian Institutes of Health Research and Canadian Institute for Veteran Health Research (to RL); and a core grant to the Waisman Center from the National Institute of Child Health and Human Development (P30-HD003352) [to DG, JN, RD].

