## Supplementary Materials for "Volumetric Differences of Thalamic Nuclei are Associated with Post-Trauma Psychopathology"

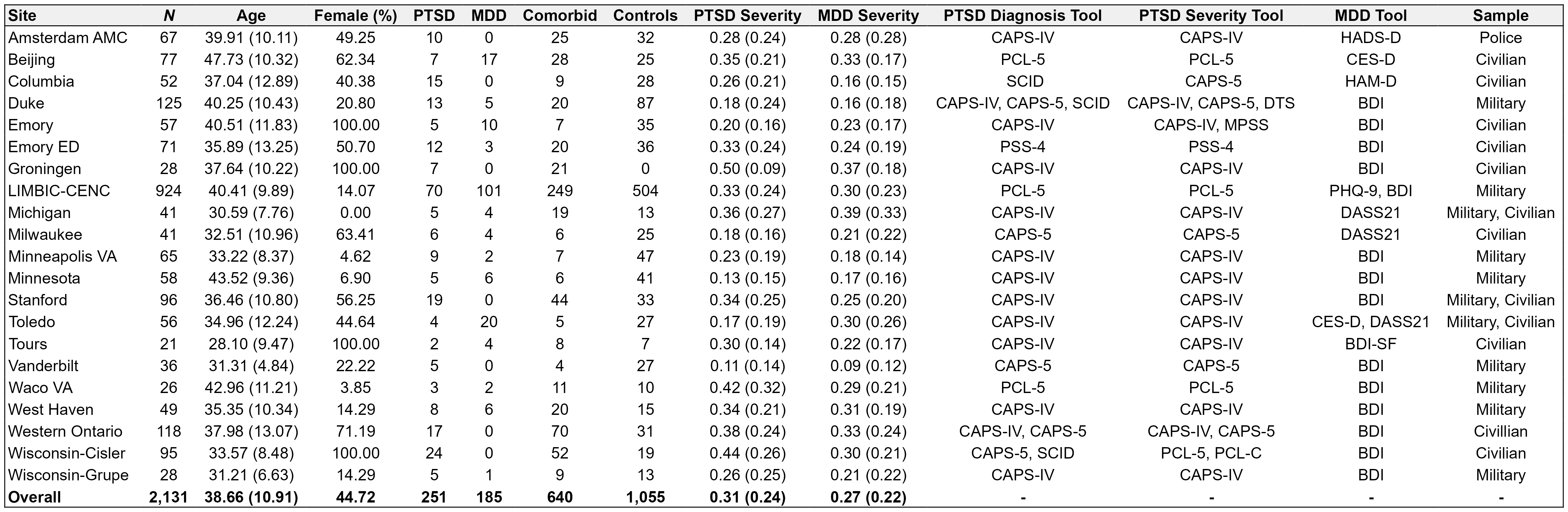

**Table S1**: Demographic information broken down by site. Means reported with standard deviations in parentheses.

| **Site** | **PI(s)** | **Inclusion criteria** | **Exclusion criteria** |
| --- | --- | --- | --- |
| Amsterdam AMC | Olff, Veltman | All: Police officers 18-65 years of age who are eligible for MRI  PTSD: current PTSD diagnosis, with CAPS ≥ 45.  Controls: exposure to at least one traumatic event (according to DSM-IV A1 criterion), with CAPS < 15 | General: History of neurological disorders, any severe or chronic systemic disease or unstable medical condition (including endocrinological disorders), use of psychotropic medications.  Females: pregnancy or breastfeeding.  PTSD: current psychotic disorder, substance-related disorder, severe personality disorder, severe major depressive disorder (MDD) (i.e., involving high suicidal risk and/or psychotic symptoms) or current suicidal risk.  Controls: any current Axis-1 disorder and lifetime history of PTSD or MDD |
| Beijing | Li | Individuals 18-65 years of age who personally experienced Wenchuan earthquake in 2008 and are right-handed | Intellectual disability; major psychosis (e.g., schizophrenia and organic mental disorders); drug or alcohol abuse; history of head trauma or surgery; metallic embedded object in body; claustrophobia; exposure to other trauma events from time of the disaster to the time of the study. |
| Columbia | Zhu, Neria | All: Males or females 18-60 years of age able to give consent, fluent in English  PTSD: Experience of a traumatic event or events during lifetime; current DSM-V Criterion A for PTSD | All: history of psychosis, bipolar disorder, or dementia; significant depression (HAM-D>25); suicidality; recent substance/alcohol dependence (past 6 months) or abuse (past 2 months); psychotropic medication usage within past 4 weeks (e.g., antipsychotics, antidepressants, mood stabilizers, or stimulants); triptan anti-migraine medications; β-blockers; pregnancy; MRI contraindications; serious and/or unstable/untreated medical illness (e.g., stroke, brain tumor, demyelinating disease)  Controls: Current or lifetime history of major psychiatric diagnosis, e.g., major depressive disorder, psychotic disorder, bipolar disorder, obsessive compulsive disorder (OCD), PTSD, panic disorder, agoraphobia, eating disorder or alcohol/substance use disorder; history of DSM-5 criterion A1 trauma exposure; HAM-D>7 |
| Duke | Morey | Veterans 18-65 years of age, fluent in English, free of implanted metal objects or metal shards in eyes | Axis I psychiatric conditions other than PTSD or MDD; current substance abuse or lifetime substance dependence (other than nicotine); high risk for suicide, claustrophobia; neurological disorders; learning disability or developmental delay; major medical conditions |
| Emory | Stevens, Fani | Individuals 18-65 years of age who speak English and have endorsed at least 1 criterion A trauma | Current psychotic symptoms or bipolar disorder; current substance or alcohol dependence; history of head trauma; psychoactive medication usage; current illegal drug use (verified with urine drug screen within 24 hours of scan) |
| Groningen | Daniels | Civilian women 20-60 years of age with a current PTSD diagnosis, sufficient proficiency in German, MRI compatible | Neurologic disorders; history of substance abuse or dependence (past 6 months); history of head injury; cerebral incidental findings verified by a neuroradiologist after the MR scan; usage of benzodiazepines, tricyclic antidepressants, or anticonvulsants; primary borderline personality disorder; other current diagnosis of Axis I disorder |
| LIMBIC-  CENC | Cifu, Walker, Wilde | Veterans with history of deployment in Operation Enduring Freedom (OEF), Operation Iraqi Freedom (OIF), Operation New Dawn (OND), or follow-up conflicts; history of combat exposure (score>1 on any item in Deployment Risk and Resiliency Inventory Section D [DRRI-2-D]) | History of moderate to severe TBI; history of major neurologic disorder with significant decrease in functional status and/or loss of ability for independent living; severe psychiatric disorder (e.g., schizophrenia) |
| Michigan | Liberzon | All: Male combat veterans and civilians who are 18-55 years of age and eligible for MRI.  PTSD: current PTSD diagnosis with CAPS-IV ≥50. Trauma-exposed controls: exposure to ≥1 traumatic event (per DSM-IV A1 criterion) with CAPS-IV <15. Community controls: CAPS-IV < 15. | History of neurological disorders, any severe or chronic disorder, current alcohol or drug abuse and/or dependence. |
| Milwaukee | Larson | Civilians aged 18-60 years; exposure to DSM-5 A1 criterion trauma; high risk for PTSD (score ≥3 OR item 2 rated ≥3 on Predicting PTSD Questionnaire, Rothbaum et al., 2014); English speaking; ability to schedule baseline study visit within 30 days of traumatic injury | Glasgow Coma Scale score ≤ 13 (i.e., moderate to severe traumatic brain injury); on police hold; contraindication to MRI; pregnancy (or planned pregnancy within 6 months); intentional self-inflicted injury; severe vision or hearing impairment; history of psychotic or manic symptoms, or neurologic condition (e.g., seizures, spinal cord injury); currently on antipsychotic medication; clear evidence of substance use disorder |
| Minneapolis VA | Disner, Davenport | OEF and/or OIF veteran 22-60 years of age who had been exposed to combat during their deployment(s). | Current psychosis; current DSM-IV substance abuse or dependence other than alcohol, caffeine, or nicotine; moderate or severe traumatic brain injury; neurologic condition other than TBI; current unstable medical condition that would likely affect brain function (e.g., uncontrolled diabetes); significant imminent risk of suicidal or homicidal behavior. |
| Minnesota | Lissek | Individuals 18-65 years of age with history of combat-related trauma | Current of past history of psychosis, bipolar disorder, delirium, dementia, amnestic disorder, or intellectual disability; suicidality; substance use disorder within past six months; pregnancy; current or past medical illnesses that may confound study results or place participant at risk; current use of any medication that alters central nervous system function including antidepressants, benzodiazepines, anti-psychotics, mood-stabilizers, anti-parkinsonian agents, anti-convulsants, sleep medications, pain medications, and anti-hypertensives; MRI contraindications |
| Stanford | Etkin, Maron-Katz | OEF/OIF Veterans 18 and 65 years of age; fluent in English; able to provide informed consent. | History of psychotic, bipolar, or substance dependence (within 3 months for PTSD group or lifetime for controls), history of neurological disorder, moderate to severe traumatic brain injury, claustrophobia, regular use of benzodiazepines, opiates, thyroid medications, or other CNS medications.  Trauma-exposed healthy controls: history of any Axis I psychiatric disorder, including PTSD. |
|  |  | All: Community-dwelling adults 18-60 years of age who are fluent in English  Patients: Individuals experiencing chronic (>3 months) moderate to severe anxiety or depression (indicated by a score >10 on the PHQ-9 [excluding suicide item] OR a score >10 on the GAD-7 scale) - must express interest in seeking treatment for psychiatric symptoms.  Controls: PHQ-9 and GAD ≤ 4. | All: contraindications to MRI or TMS; current participation in psychiatric treatment; history of neurological disorder, brain surgery, electroconvulsive or radiation treatment, brain hemorrhage or tumor, stroke, epilepsy, hypo- or hyper-thyroidism; medication use that substantially reduces seizure threshold to TMS (e.g., olanzapine, chlorpromazine, lithium) and unwillingness or inability to safely withdraw at least two weeks prior to TMS appointment; medications that interfere with blood flow (e.g., opiates, antihypertensive medication); insufficiently controlled thyroid dysfunction; current substance dependence (within past 3 months); refusal to abstain from illicit drug use for study duration; refusal to abstain from alcohol within 24 hours of MRI scans; pregnancy; prior exposure to deep brain stimulation, rTMS, or tDCS therapies; significant traumatic brain injury (indicated by loss of consciousness, post-trauma amnesia, imaging findings, penetrating brain injury); history of psychotic or manic symptoms. |
| Toledo | Wang | Motor vehicle accident (MVA) survivors transported to the University of Toledo Emergency department, or to a ProMedica emergency medicine department. | Pregnancy; under the influence of alcohol or drugs at the time of MVA; major injuries, moderate to severe traumatic brain injury; major medical illnesses; contraindication to MRI |
|  |  | Ohio National Guard and Reserve soldiers 18-50 years of age who were deployed in OEF or OIF – must have met Ohio National Guard Study characteristics and able to provide informed consent. | History of psychosis, bipolar disorder, or neurologic condition; current substance dependence; intellectual disability or developmental disorder; contraindication to MRI; current use of antipsychotic medication. |
| Tours | Quidé, El Hage | Female survivors of sexual assault (within past month)  Control: no history of interpersonal violence exposure | History of head injury, substance abuse, current use of psychotropic medication, medical conditions with confounding effects on brain function (e.g., epilepsy, brain tumor), contraindication to MRI |
| Vanderbilt | Blackford | OEF/OIF/OND Veterans 18-50 years of age who are fluent in English | Psychoactive medication usage in past 6 weeks; participated in psychotherapy within the past month; current substance use disorder (>6 month remission); positive urine drug or alcohol breath screen on MRI study day; history of psychotic or bipolar disorder, traumatic brain injury, or significant medical (e.g., cancer, HIV) or neurological illness (e.g., stroke, brain tumor, multiple sclerosis, epilepsy); contraindication to MRI  Trauma-exposed controls: lifetime diagnosis of PTSD; symptoms of hypervigilance  Healthy controls: any trauma exposure |
| Waco VA | May, Gordon | Veterans 18-60 years of age | Seizure disorder, dementia; MRI contraindications |
|  |  | Veterans 18-60 years of age with a clinical diagnosis of TBI in their VA medical record | Diagnosis of schizophrenia, schizoaffective disorder, bipolar disorder type I, severe substance use disorder, or a high risk of suicide; absence of qEEG more than 2SDs outside of population means of healthy age-matched controls; MRI contraindications |
| West Haven | Abdallah | Combat-exposed US Veterans aged 21-65 both with and without PTSD, fluent in English | History of psychotic disorder, bipolar depression, or neurologic/neurodevelopmental disorder (including learning disorders, attention-deficit hyperactivity disorder, moderate to severe traumatic brain injury, epilepsy, brain tumor); contraindication to MRI |
| Western Ontario | Lanius | Primary diagnosis of PTSD for patients | Incompatibilities with scanning conditions, previous neurologic and development illness, comorbid schizophrenia or bipolar disorder, alcohol or substance abuse, a history of head trauma, or pregnancy during scan. participants were excluded if they had implants or metal that do not comply with 3T fMRI safety standards for research, a history of head injury with a loss of consciousness, significant untreated medical illness, a history of neurological disorders, history of any pervasive developmental disorders, pregnancy, and current use of any psychotropic medication within one month prior to study. PTSD individuals were further excluded if they reported a history of bipolar disorder, schizophrenia, or substance-use disorder prior to participation of the study |
| Wisconsin-Cisler | Cisler | Women aged 21-50 with and without history of interpersonal violence – must be fluent in English | History of psychosis, medication changes within past 4 weeks, cognitive impairment, current substance or alcohol use disorder |
| Wisconsin-Grupe | Grupe | Adults 18-50 with exposure to 1+ life-threatening war zone trauma events; capable of giving informed consent and fluent in English; clear evidence of war zone trauma exposure in Iraq or Afghanistan since 2001 (e.g., Combat Action Ribbon [Marines], Combat Infantry Badge [Army]); stable pharmacological or psychotherapeutic treatment for at least 8 weeks prior to beginning of study | Weight >352 pounds or over; pregnancy or current breastfeeding; Metallic implants such as prostheses or aneurysm clip, or electronic implants such as cardiac pacemakers; Neurological or serious medical condition; History of seizures or seizure disorder; Moderate or severe traumatic brain injury; Current active substance dependence or dependence within 3 months (other than nicotine); bipolar disorder, schizophrenia, schizoaffective disorder, psychotic disorder NOS, delirium, or any DSM-IV cognitive disorder; Severe psychiatric instability or severe situational life crises, (e.g., suicidality, homicidality); extensive experience in yoga or meditation; Current use of benzodiazepines and beta-blockers |

**Table S2**: Inclusion and exclusion criteria for each site.

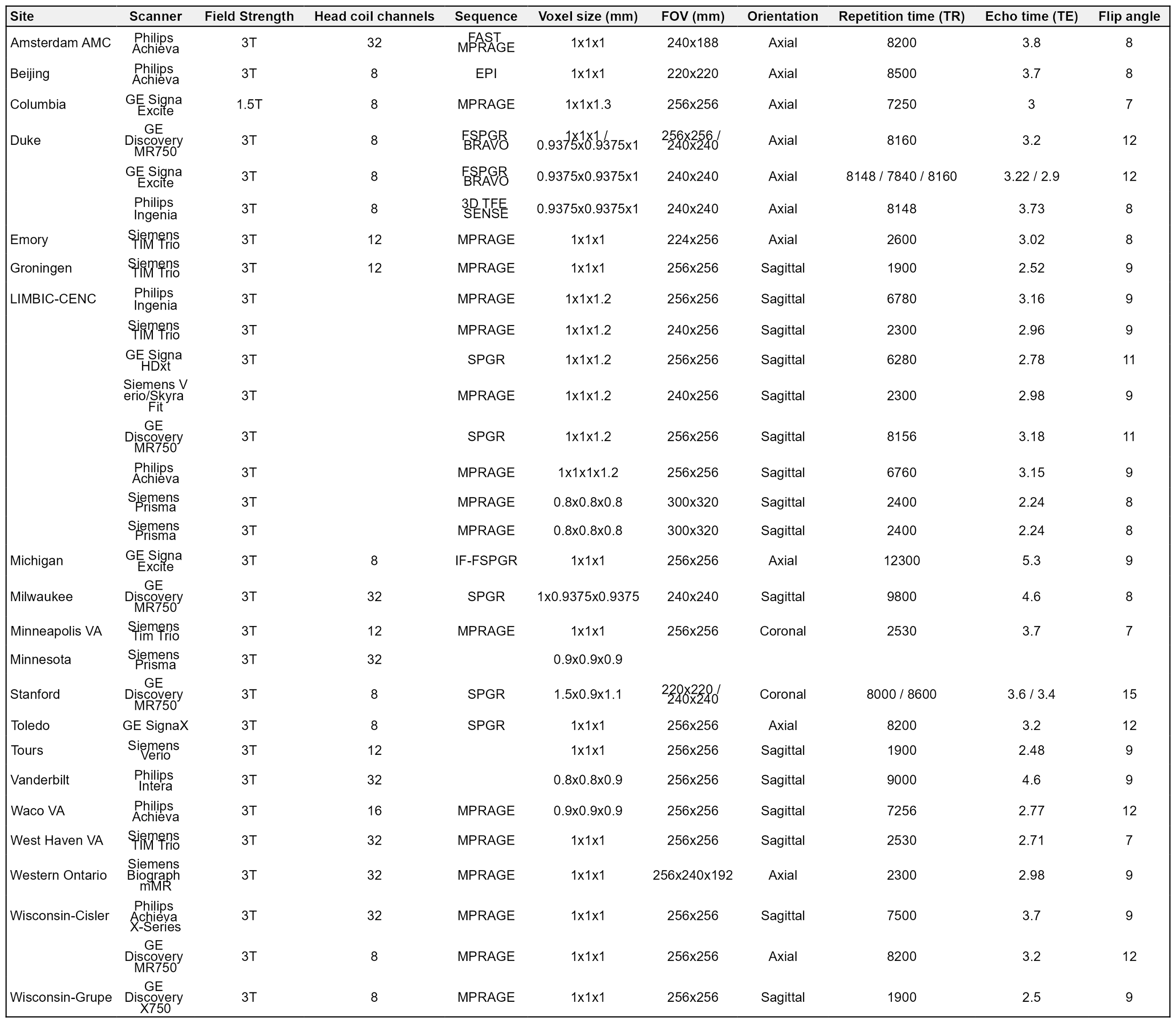

**Table S3**: MRI scanning parameters of each site.

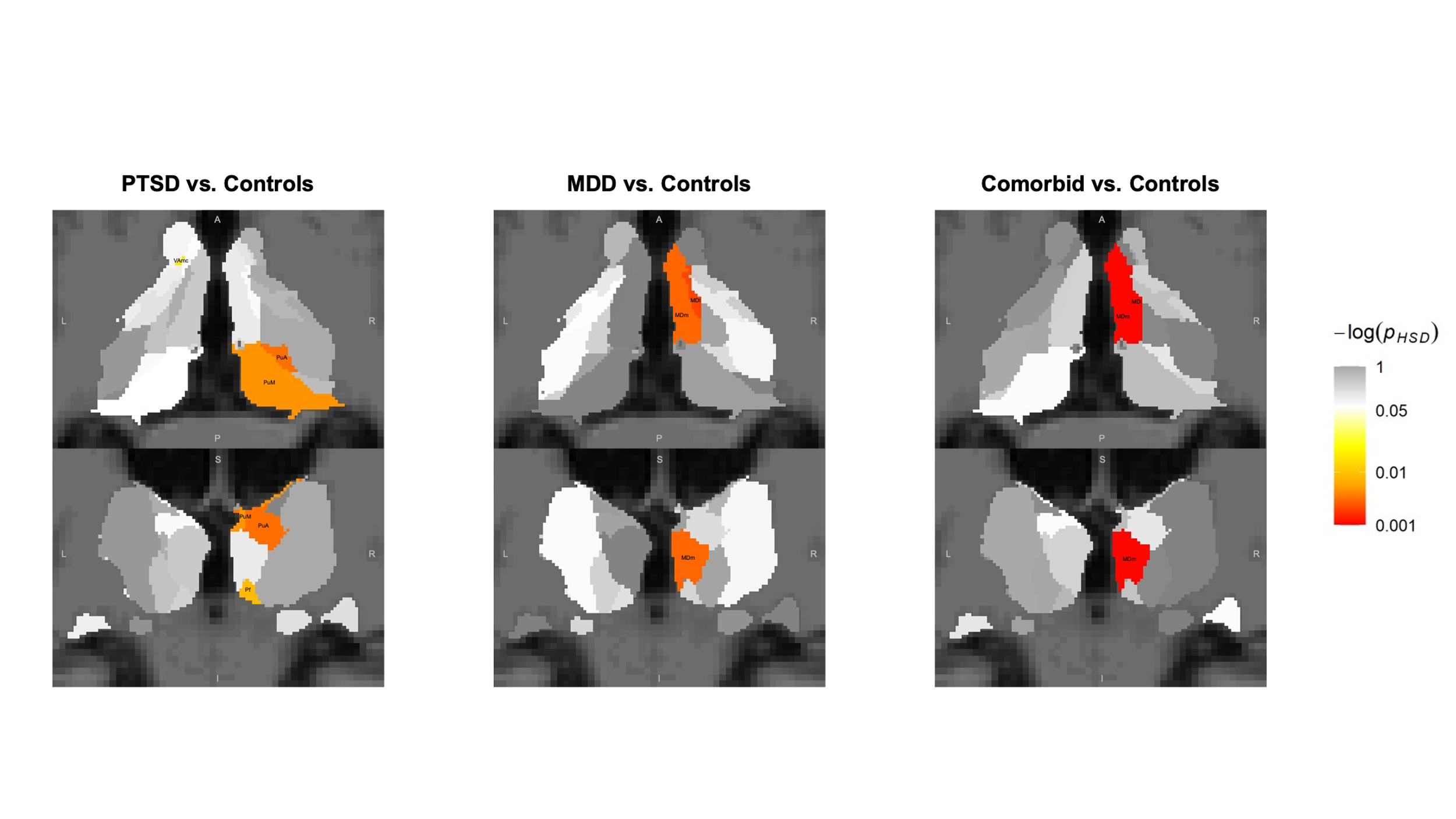

**Figure S1**: Heatmap of the thalamus displaying HSD-corrected *p*-values of the diagnostic groups compared to controls. Values in the legend have been log-transformed.

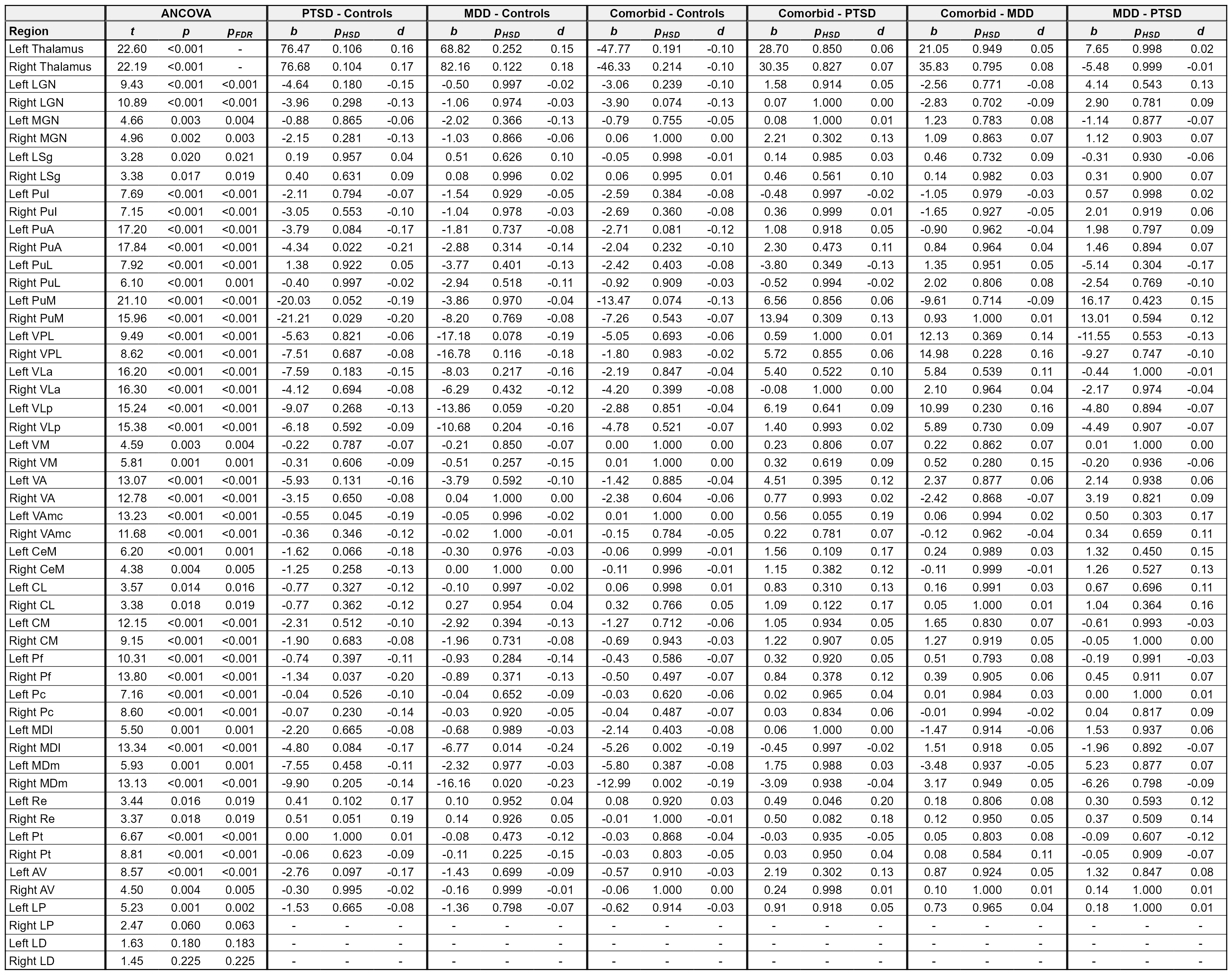

**Table S4**: ANCOVAs were performed comparing diagnostic groups (Equation 1) with post-hoc pairwise comparisons of significant ANCOVA results. ANCOVA *p*-values have been FDR-corrected for 50 comparisons. Pairwise *p*-values have been corrected using the Tukey HSD method to correct for the six group comparisons.
